# Ancient U/V sex chromosomes reveal slow expansion of recombination suppression in Polytrichaceae mosses

**DOI:** 10.64898/2026.10.30.763134

**Authors:** Fabien Rey-Giraud, Muskaan, Sara Oort, Alain Vanderpoorten, Nils Cronberg, Christelle Blassiau, Isabelle Charissou, Xavier Vekemans, Christelle Fraïsse

## Abstract

Sex chromosomes evolving in haploid versus diploid life cycles provide complementary systems for testing theories of sex chromosome evolution, yet haploid sex chromosomes remain poorly studied. In dioicous bryophytes, females and males carry non-recombining U and V chromosomes, respectively, providing a unique opportunity to examine the consequences of long-term recombination suppression independently of diploid heterogamety. Here, we characterize the U/V sex chromosomes of Polytrichaceae using chromosome-scale genomes from ten species. We identify homologous U/V chromosomes across the family and show that their non-recombining sex-determining regions (SDRs) originated early in Polytrichaceae evolution, approximately 124 million years ago. Despite their ancient origin, however, the SDRs have undergone limited expansion and are consistently flanked by large pseudoautosomal regions (PARs). The SDRs nevertheless show strong genomic signatures of recombination suppression, including reduced gene density, transposable-element accumulation, reduced diversity, high synonymous divergence, and altered codon usage. Yet they retain hundreds of genes and show limited evidence of functional degeneration. Their gene content is dynamic, with frequent lineage-specific gains and losses, while a conserved core of 60 orthogroups has persisted across all ten species for more than 100 million years. Comparisons across sex chromosome systems suggest that haploid U/V chromosomes retain more genes than similarly old diploid X/Y or Z/W chromosomes, consistent with a role for haploid selection in limiting degeneration. However, haploid selection alone cannot explain the limited SDR expansion observed in Polytrichaceae, suggesting that other factors also shape SDR evolution. Together, these results reveal an ancient U/V system in which recombination suppression has strong genomic consequences but has not driven extensive SDR expansion or functional degeneration. By decoupling recombination suppression from diploid heterogamety, U/V chromosomes provide a powerful framework for distinguishing the consequences of recombination arrest from those of ploidy in sex chromosome evolution.

**Teaser text:** Sex chromosomes are often found to degenerate as recombination is progressively lost. But what happens when recombination stops in a haploid organism? We investigated this question in Polytrichaceae mosses, an understudied lineage with ancient U and V sex chromosomes. Across ten species, we show that these chromosomes have been non-recombining for more than 100 million years and bear strong genomic signatures of recombination suppression. Yet their non-recombining regions remain remarkably small but retain hundreds of functional genes, while large pseudoautosomal regions persist. These findings suggest that recombination suppression and sex chromosome degeneration are not inevitably coupled. Haploid selection may contribute to the retention of functional genes, but cannot alone explain the limited expansion of the non-recombining region. By providing a natural contrast to diploid XY or ZW sex chromosomes, ancient U and V chromosomes offer a powerful system for disentangling the effects of recombination suppression from those of diploid heterogamety.

## Introduction

Understanding sex chromosome evolution is central to evolutionary biology because it provides insights into how suppressed recombination, reduced effective population size, and sex-linked inheritance shape genome architecture. Classical theory posits that sex chromosomes originate through the suppression of recombination around a sex-determining locus, forming a sex-determining region (SDR). In diploid organisms, the XY and ZW sex chromosome systems are typically characterized by degeneration of the non-recombining chromosome (Y/W) in the heterogametic sex (i.e., XY males or ZW females), due to processes such as Muller’s ratchet, genetic hitchhiking, and background selection (Sigeman et al., 2024; Bachtrog, 2013; B. Charlesworth, 1978). Why recombination becomes suppressed beyond the genes directly involved in sex determination remains an open question. Numerous theoretical models have been proposed to explain SDR expansion in XY and ZW systems despite the long-term costs associated with suppressed recombination (Jay et al., 2024; Saunders & Muyle, 2024). Sexually antagonistic selection has traditionally been considered the primary mechanism driving SDR expansion. More recent models propose that the accumulation and sheltering of recessive deleterious mutations on the SDR of the heterogametic sex can also favor the expansion of suppressed recombination (Jay et al., 2026; Lenormand & Roze, 2022; Olito & Abbott, 2025); reviewed in B. Charlesworth & Olito, 2024. However, these theoretical predictions are largely derived from diploid sex chromosome systems, neglecting the role of haploid selection.

Although flowering plants are predominantly hermaphroditic, dioecy has evolved repeatedly and is often associated with sex chromosomes. Most characterized plant sex chromosome systems have a relatively recent origin compared with those described in animals (Renner & Müller, 2021). Nevertheless, plants show a high prevalence of homomorphic sex chromosomes, and some studies showed that the heterogametic chromosome had retained a large fraction of its ancestral gene content. When comparing sex chromosome systems of similar ages, plants also lack the extreme Y/W degeneration observed in some animals (Rousselle & Pannell, 2026). This reduced degeneration may reflect selection acting during the haploid phase (e.g. pollen tube growth).

Although most sex chromosome research has focused on organisms with predominantly diploid life cycles, many eukaryotes, including algae and bryophytes, have haplontic or haplodiplontic life cycles in which the haploid phase bears the sexual organs (Božović et al., 2024). In these systems, sex chromosomes are typically designated U and V in females and males, respectively. Because the diploid phase is short-lived and non-sexual, recombination between U and V is completely suppressed, exposing both chromosomes to the genomic consequences of recombination arrest. Accordingly, U and V chromosomes often show increased repeat accumulation and reduced gene content (Barrera-Redondo et al., 2025; Carey et al., 2021). However, sex-specific genes expressed during the haploid phase remain directly exposed to selection, which is expected to constrain their loss from U and V chromosomes (Bull, 1978; D. Charlesworth, 2022). Despite this prediction, direct empirical evidence for reduced degeneration driven by haploid selection remains limited. Some of the oldest sex chromosomes known do occur in organisms with U/V systems, including chromosomes estimated to be >400 Mya in Marchantiales and potentially in Ochrophyta (Barrera-Redondo et al., 2025; Bowman et al., 2017), consistent with long-term retention of gene content. In mosses, however, our knowledge about U/V chromosome evolution is restricted to few independent systems belonging to the Bryopsida and Sphagnopsida lineages (Gao et al., 2023; Healey et al., 2023).

Here, we provide the first comparative molecular characterization of U/V sex chromosomes in the early-diverging moss class Polytrichopsida. We investigate their evolutionary history by comparing all ten Polytrichaceae species with available chromosome-scale genome assemblies. We first delineate the sex chromosomes using genomic data from sexed individuals before assessing their degeneration and patterns of gene gain and loss across the family, providing the first comparative view of U/V chromosome evolution in this lineage. We show that Polytrichaceae SDRs have undergone limited expansion despite their ancient origin, and to our knowledge, it also represents the first characterized SDR/PAR system in mosses.

## Methods

**Table S1** summarizes the datasets used in this study, the main associated analyses, and their objectives, providing an overview of the study’s methodological framework.

### Plant material and reference genomes

We included eleven chromosome-scale genome assemblies available from NCBI, comprising ten Polytrichaceae species spanning five genera and the outgroup *Tetraphis pellucida* (Tetraphidaceae). Species names and assembly information are provided in **Table S2**. We sampled gametophytes from natural populations of two Polytrichaceae species, *Polytrichum commune var. perigoniale* and *Pogonatum aloides*. For each species, we collected four male and four female gametophytes for whole-genome shotgun sequencing, as well as 24 female gametophyte–sporophyte pairs for transcriptome sequencing. We additionally collected 24 unsexed gametophytes for whole-genome shotgun sequencing in *P. commune var. perigoniale* and in *P. commune var. commune*, as well as two unsexed gametophytes of *Polytrichum juniperinum* (used as an outgroup). The four samples initially identified as female *P. commune var. perigoniale* were subsequently found to be *Polytrichum formosum*. Sample information is provided in **Table S3**.

### DNA and RNA sequencing

For DNA and RNA sequencing, whole samples were flash-frozen in liquid nitrogen and homogenized using ceramic beads. Genomic DNA was extracted using the NucleoSpin Plant II Mini Kit, and total RNA using the RNA PLUS XS Kit (Macherey-Nagel, Düren, Germany). DNA and RNA concentrations were measured using a Qubit 2.0 fluorometer (Thermo Fisher Scientific). RNA quality was assessed using a Fragment Analyzer (FA5200, Agilent Technologies) with the DNF-471 kit. DNA libraries were prepared using the NEXTFLEX Rapid XP V2 DNA-Seq Kit, and RNA libraries using the NEXTFLEX Rapid Directional RNA-Seq Kit 2.0, with NEXTFLEX Poly(A) Beads for poly(A) RNA selection. Library concentrations were measured using a Qubit 2.0 fluorometer with the DNA HS Assay Kit, and library quality was assessed using a Fragment Analyzer with the DNF-473 kit. Libraries passing quality control were sequenced by the Genotoul GeT-PlaGe platform (INRAE, Castanet-Tolosan, France) to generate 150-bp paired-end reads on an Illumina NovaSeq S4 300 platform (Illumina, San Diego, CA, USA). The DNA samples from sexed *P. commune var. perigoniale* individuals were instead sequenced on an AVITI3 platform (Element Biosciences, San Diego, CA, USA).

### Genome annotation

Transposable elements (TEs) in the eleven genomes were annotated and soft-masked using EarlGrey v4.4.5 (Baril et al., 2024). For *P. commune* and *P. aloides*, RNA-seq reads were mapped to the respective reference genomes using HISAT2 v2.2.1 (Kim et al., 2019) and assembled into transcriptomes with StringTie v2.2.1 (Kovaka et al., 2019). Coding sequences were predicted from the assembled transcripts using TransDecoder v5.5.0 (Haas & Papanicolaou, 2015), generating species-specific protein sets. These proteins were combined with reference proteins from the BUSCO bryophyta_odb10 dataset (Zhou et al., 2025) to construct a Polytrichaceae protein database. This database was used to annotate all eleven genomes with Galba v1.0.11 (Brůna et al., 2023). Annotation completeness was assessed using BUSCO v5.7.1 (Simão et al., 2015) with the bryophyta_odb10 dataset, and results were summarized using MultiQC v1.17 (Ewels et al., 2016; **Figure S1**).

### Read cleaning and variant calling

Raw DNA reads were quality-filtered using fastp v0.23.4 (S. Chen, 2023) and screened for potential contaminants using a custom Kraken2 v2.1.3 database (Wood et al., 2019) containing UniVec_Core sequences, fungi, bacteria, archaea, viruses, plasmids, protozoa, and human sequences. Reads passing these filters were aligned to the corresponding reference genome using BWA v0.7.17 (Li, 2013). Variants were called with GATK v4.5.0.0 (Poplin et al., 2018) using HaplotypeCaller in haploid mode (--sample-ploidy 1, -ERC GVCF, --min-base-quality-score 30, -stand-call-conf 30). GVCFs from samples of the same population or sex were subsequently combined using CombineGVCFs, followed by joint genotyping with GenotypeGVCFs (-ploidy 1, -all-sites TRUE). Before calculating SNP density and sequencing depth along the genome, TE-associated positions were removed from the VCFs generated from sexed *P. commune var. perigoniale* and *P. aloides* samples to minimize biases in SNP density and depth estimates arising from repetitive regions. SNPs were filtered using GATK VariantFiltration based on site quality (QUAL > 30), genotype quality (GQ > 30), and depth of coverage (DP > 5). Population genetic analyses were performed using the unsexed *P. commune var. perigoniale* samples. For these analyses, VCFs were additionally filtered to retain biallelic SNPs, remove indels, and exclude sites with >20% missing data using samtools v1.16.1 (Danecek et al., 2021).

### Gene expression

We used the *de novo* transcriptomes of *P. commune* var. *perigoniale* and *P. aloides* generated by Muskaan et al. (in prep.). RNA-seq reads were pseudo-aligned to the corresponding *de novo* transcriptomes using Salmon v1.10.2 (Patro et al., 2017) to obtain transcript-level read counts. Gene expression levels were then quantified by averaging the log2-transformed expression values obtained with edgeR v4.2.2 (Y. Chen et al., 2025) and limma v3.60.6 (Ritchie et al., 2015). Finally, we used the `filterByExpr` function in edgeR to identify genes with sufficient expression support and removed genes that did not meet this criterion, which were considered potential pseudogenes.

For molecular evolution and codon usage analyses, coding sequences (CDS) from the genome annotations were compared with CDS supported by expression in the *de novo* transcriptomes using BLAST v2.17.0 (Camacho et al., 2009), with an E-value threshold of 1e-15 and retaining only the best hit. We retained annotated CDS with a BLAST hit covering ≥75% of their length at ≥90% sequence similarity in at least one of the two transcriptomes. This filtering step reduced the contribution of poorly supported or potentially pseudogenized genes to downstream analyses.

### Identification of sex chromosomes

Sex-determining regions (SDRs) were identified in *P. commune* and *P. aloides* by scanning the genome for differences in SNP density and read depth between male and female samples using VCFs generated from sexed individuals. Read depth was calculated in genomic windows of 1Mb and normalized to the genome-wide mean for each sample. We defined the SDR boundaries as the genomic positions where non-sex-specific patterns transitioned to sex-specific patterns in read depth and SNP density. The PAR showed similar read depth and SNP density in males and females, whereas the SDR showed a reduced male-to-female read-depth ratio and elevated male SNP density. Boundaries were placed at the transition between these contrasting patterns. In both species, the SDRs were characterized by higher GC content, higher long-terminal repeat density, and lower gene density than the autosomal regions. These genomic signatures were then used to delineate putative SDRs in the remaining Polytrichaceae species by placing their boundaries at the points where genome-wide trends shift toward locally elevated or reduced feature values. No comparable patterns were detected in *T. pellucida*, consistent with the absence of sex chromosomes in this monoicous species. Correspondence among putative SDRs across Polytrichaceae genomes was assessed using whole-genome synteny comparisons with jcvi v1.5.12 (Tang et al., 2024).

### Phylogenetic reconstruction

A species tree was inferred from 139 BUSCO genes (bryophyta_odb10) identified as single-copy orthologs in all genomes, to reconstruct the evolutionary relationships among the studied Polytrichaceae species. Coding sequences for each gene were aligned across species using MACSE v1.2 (Ranwez et al., 2011). Poorly aligned blocks were trimmed with BMGE v1.12 (Criscuolo & Gribaldo, 2010), concatenated using AMAS v1.0 (Borowiec, 2016), and subsequently filtered for outlier sequence fragments using spruceup v2024.7.20.post1 (Borowiec, 2019). Maximum-likelihood phylogenetic inference was performed using IQ-TREE v3.0.1 (Wong et al., 2026) using ModelFinder Plus to select and merge substitution models across BUSCO genes (-m MFP+MERGE -merit AICc). Branch support was assessed using 1,000 ultrafast bootstrap replicates with NNI optimization and 1,000 SH-aLRT tests, with -rclusterf 4 used to reduce computational time.

### Time calibration

Divergence times were estimated using BEAST v2.7.7 (Bouckaert et al., 2019) under a GTR substitution model and a strict molecular clock. The divergence between Polytrichaceae and *T. pellucida* was calibrated using the estimate of 337 Mya from Bechteler et al. (2023), implemented as an offset normal prior with an onset of 337 Ma, mean = 0 and sigma = 5. This calibration represents an estimated divergence time, with uncertainty around the published estimate. Fossil evidence from the genus Atrichum, dated to 37–57 Mya (Frahm, 2004), was used to calibrate the divergence between *Atrichum angustatum* and *Oligotrichum hercynicum*. Because this fossil provides a minimum age constraint rather than a point estimate of the divergence time, this calibration was implemented as a gamma prior with an offset of 37 Ma, alpha = 2, and beta = 10.

### Molecular evolution analyses

Polymorphism within *P. commune var. perigoniale* and divergence from *Polytrichum juniperinum* were estimated using dNdSpNpS v3.0 (Loire et al., 2013) on the dataset filtered for potential pseudogenes (see “Gene expression”). Default options were used, except that sites with genotype calls in fewer than three individuals (*gapN_site*) were excluded. For biallelic sites, we estimated the numbers of synonymous (S) and nonsynonymous (N) sites, as well as synonymous (πS) and nonsynonymous (πN) polymorphism within *P. commune var. perigoniale*. Synonymous (dS) and nonsynonymous (dN) divergence were estimated between *P. commune var. perigoniale* and *P. juniperinum*. Confidence intervals were calculated from 10,000 bootstrap replicates over genes.

### SDR tree reconstruction

Gene families within the SDRs of the ten Polytrichaceae species were identified by clustering their predicted proteins using OrthoFinder v3.1.3 (Emms et al., 2026). We identified 50 single-copy gene families shared across all SDRs and used them to infer an SDR gene tree, including sexed haplotypes from *P. aloides* and *P. commune var. perigoniale*. The gene tree was inferred following the procedure described in the “Phylogenetic reconstruction” section. Only 40 gene families were retained after filtering by BMGE v1.12 (Criscuolo & Gribaldo, 2010). Gene gains and losses across the SDR phylogeny were inferred using GLOOME v1.266 (Cohen et al., 2010).

### Estimation of synonymous divergence

Synonymous divergence among species (dS) and between U/V gametologs (ks) were estimated using dNdSpNpS v1.0 using default parameters except for gapN_site which is set to allow sites supported by a single individual. For each of the 139 BUSCO gene used in the species phylogeny, pairwise dS values were calculated between species of the Atrichum, Oligotrichum, and Polytrichastrum genera and species of the Pogonatum and Polytrichum genera, providing estimates of synonymous divergence associated with the Polytrichaceae crown age. Synonymous divergence between U and V gametologs was estimated for sexed *P. aloides* and *P. commune* samples using the same pseudogene filtering criteria described in “Gene expression”. Only single-copy SDR genes inferred to be ancestral based on the GLOOME analysis were retained.

### Estimation of SDR age

Using a resampling approach, we estimated SDR age from neutral synonymous divergence, assuming that synonymous divergence accumulates approximately linearly over time. The Polytrichaceae crown age (**Figure S2**), its corresponding dS distribution (**Figure S3**), and the Ks distribution of SDR gametologs (**Figure 2c**) were each modeled as independent normal distributions. For 10,000 iterations, one value was randomly drawn from each distribution, and SDR age was estimated by proportional scaling of SDR gametolog divergence relative to synonymous divergence at the Polytrichaceae crown node. The resulting distribution of SDR age estimates was summarized by its median and 95% interval. Because the SDR is shared across all Polytrichaceae species and therefore predates their diversification, the lower bound of the SDR age distribution was constrained to the lower limit of the 95% highest posterior density (HPD) interval of the Polytrichaceae crown age estimated with BEAST.

### Codon usage

Codon usage bias was compared among SDR, PAR, and autosomal regions across all Polytrichaceae species using the R package cubar v1.2.0 (Liu et al., 2026). Analyses were performed on the dataset filtered for potential pseudogenes (see “Gene expression”), excluding genes shorter than 200 codons. Optimal codons were defined as the codons preferentially used in the top 10% most highly expressed genes in the *de novo* transcriptomes of *P. aloides* and *P. commune var. perigoniale*. For each genomic partition, we calculated GC content at the third codon position (GC3s) and the fraction of optimal codons (FOP). Statistical comparisons among conspecific partitions were performed using pairwise Kolmogorov–Smirnov tests implemented in the R package Mtest v1.0.4 (Oñate & Morales-Oñate, 2023).

## Results

### Polytrichaceae U/V chromosomes display typical features of moss sex chromosomes but an unusual SDR/PAR organization

In *Pogonatum aloides*, the beginning of chromosome 4 comprises a ∼11-Mb region (∼18% of the chromosome) characterized by elevated SNP density in males, whereas SNP density in females is similar to that of autosomes (**Figure 1**). The male-to-female read depth ratio was also markedly reduced in this region, consistent with a sex-determining region (SDR) on chromosome 4 and a female reference genome. Similar patterns were observed on chromosome 4 in *Polytrichum commune* (**Figure S4**), although the contrast between males and females was weaker because the male samples were identified as belonging to *P. commune var. perigoniale*, closely related to the reference genome (*P. commune* var. *commune*) whereas the females were erroneously sampled from the more distantly related *Polytrichum formosum* (**Figure S5**). The *P. commune* SDR spans ∼17 Mb, corresponding to ∼29% of the chromosome.

**Figure 1.**
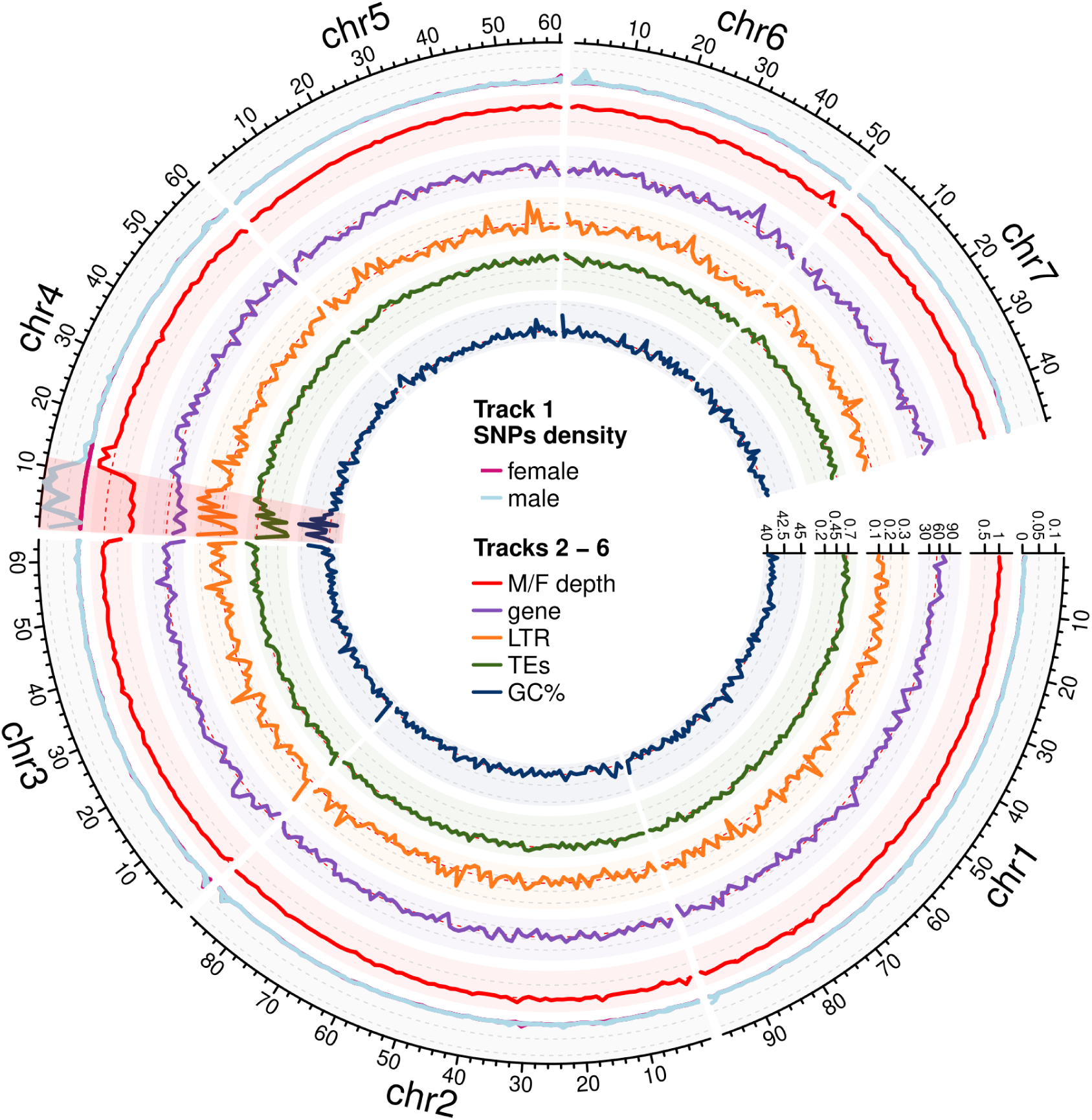
The haploid sex chromosome of *Pogonatum aloides* exhibits an SDR/PAR structure. Genomic features are shown in 1-Mb sliding windows. From top to bottom: (1) SNP density in male (blue) and female (pink) samples; (2) male over female sequencing depth ratio; (3) number of genes; (4) long-terminal repeat transposable element density; (5) total transposable element density; and (6) GC content. The SDR on chromosome 4 is shaded in red.

As observed in other bryophyte sex chromosomes (**Table S4**), including *Polytrichastrum alpinum* (Zeng et al., 2025), the SDR of *Pogonatum aloides* (**Figure 1**) and *P. commune* (**Figure S4**) as well as eight other Polytrichaceae species (**Table S5**) showed reduced gene density (∼0.38-fold that of autosomes) and increased long-terminal repeat density (∼2.7-fold that of autosomes), together with elevated GC content. The pattern of accumulation of repetitive elements in these regions is consistent with long-term suppression of recombination on the U and V chromosomes. Based on these genomic features, we identified putative SDRs ranging from ∼14 to 22 Mb in all other Polytrichaceae genomes (**Figure S6**). No SDR was detected in *Tetraphis pellucida*, consistent with its monoicous reproductive system. In *Atrichum angustatum*, by contrast, two putative SDRs corresponding to the U and V chromosomes were detected, likely resulting from the inadvertent inclusion of both a male and a female individual in the sequencing dataset (**Table S6**).

Unlike the moss sex chromosomes described to date (**Table S4**), Polytrichaceae SDRs were consistently associated with a large pseudo-autosomal region (PAR), spanning ∼66–82% of the U/V chromosome. PARs showed gene density, TE density, and GC content similar to those of autosomes (**Figure S6**). In all Polytrichum species, the PAR accounted for ∼71% of the sex chromosome. In *A. angustatum* and *P. alpinum*, the SDR (including U and V SDRs for *A. angustatum*) and PAR were located on two separate scaffolds, probably reflecting assembly fragmentation associated with Oxford Nanopore sequencing. The absence of microchromosomes in these species in cytological studies supports this interpretation (Patel et al., 2021). A similar assembly-related separation of sex chromosome regions has been reported for the U chromosome of the liverwort *Blasia pusilla*, which is enriched in tandem repeats (Yue et al., 2026).

### Polytrichaceae U/V chromosomes have a unique and ancient origin

Macrosynteny analysis revealed high collinearity among Polytrichaceae genomes (**Figure 2a**), consistent with previous observations in mosses (Kirbis et al., 2025). Notably, both the SDR and PAR were homologous across all studied species. The regions identified as putative SDRs and PARs based on their genomic features corresponded closely to the syntenic blocks defined using male–female genome comparisons in *P. aloides* (**Figure 2a**, **Table S7**). The one-to-two syntenic correspondence observed for *Polytrichastrum sexangulare* reflects its recent polyploid origin (Derda & Wyatt, 2000), with 14 chromosomes compared with seven in the other species. Microsynteny analysis between the U and V chromosomes of *A. angustatum* revealed extensive gene reshuffling, consistent with long-term recombination suppression (**Figure S9**).

**Figure 2.**
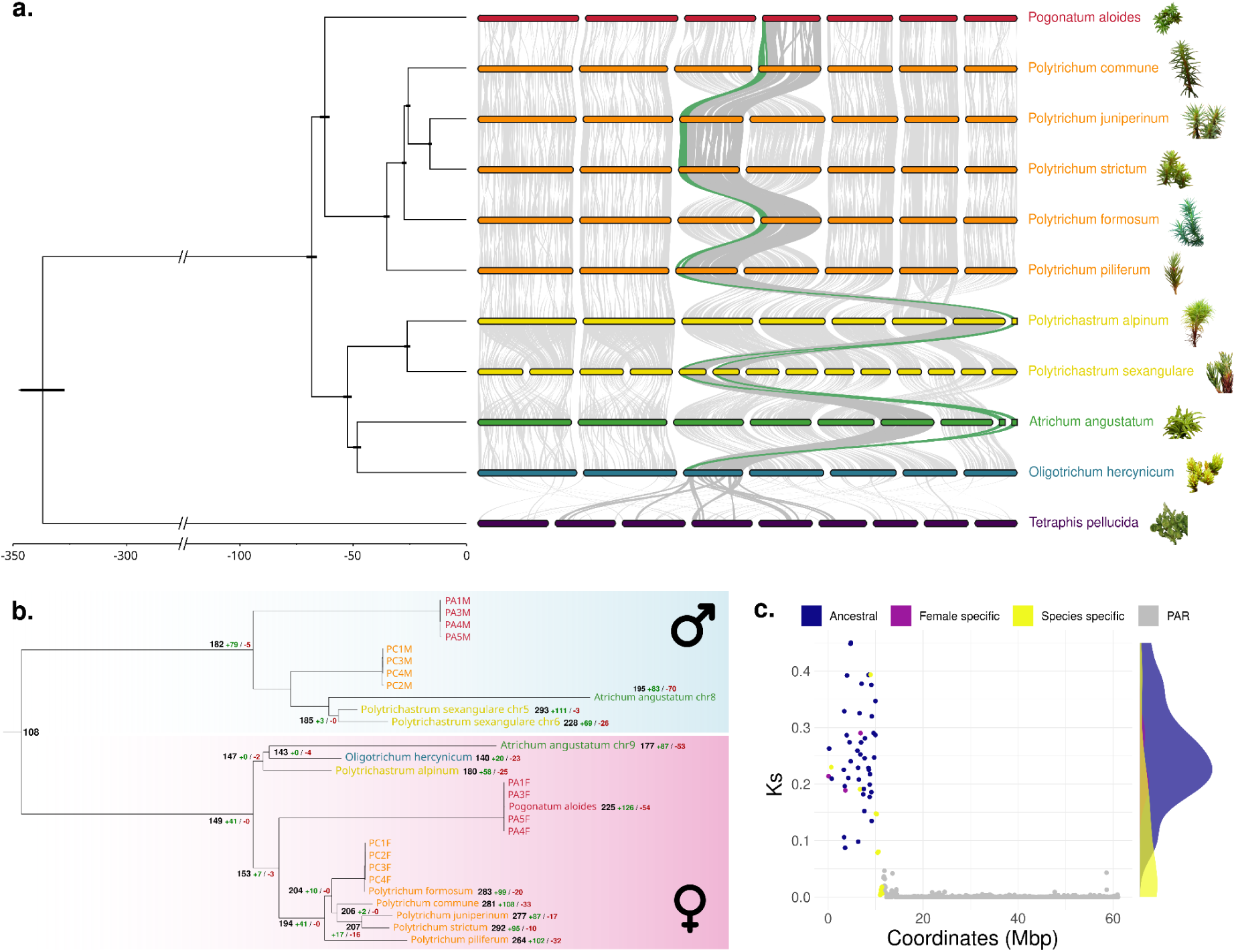
The U/V sex chromosomes of Polytrichaceae have a unique and ancient origin. **(a)** Time-calibrated BEAST phylogeny of the studied Polytrichaceae species inferred from 139 BUSCO genes, with genome-wide macrosyntenic blocks correspondence. The full BEAST tree and the IQ-TREE topology with bootstrap support are shown in **Figures S7** and **S8**, respectively. The mean Polytrichaceae crown age was estimated at 68 Mya (**Figure S2**). Colors indicate genera. Light grey, dark grey, and green ribbons indicate macrosyntenic blocks corresponding to autosomes, PAR, and SDR regions, respectively, based on their direct or indirect syntenic relationships with *Pogonatum aloides*. Within each species, chromosomes are arranged in ascending numerical order according to their published reference assemblies. **(b)** Evolutionary history of U- and V-linked SDR haplotypes inferred from 40 orthogroups shared across all studied species, with a single SDR copy in each species. Colors indicate genera. U- and V-linked haplotypes form two divergent clades. Gene gains (green) and losses (red) are indicated across the species phylogeny, excluding sexed samples; together with inferred gene numbers at internal nodes and tips. Sexed samples are abbreviated as follows: PA-M, *P. aloides* male; PA-F, *P. aloides* female; PC-M, *Polytrichum commune var. perigoniale* male; and PC-F, *Polytrichum formosum* female. **(c)** Ks divergence between U- and V-linked gametologs in *P. aloides*, plotted along the sex chromosome. Genes are colored according to their genomic location: PAR (grey), *P. aloides*-specific SDR (yellow), U-specific SDR genes (purple), and ancestral SDR genes shared across all Polytrichaceae species (blue). Distributions of Ks values for ancestral and species-specific gametologs are shown on the right.

To reconstruct the evolutionary history of the SDR, we inferred a phylogeny from 40 single-copy gametologs conserved across all Polytrichaceae species (**Figure 2b**). U and V haplotypes formed two distinct clades, within which the species phylogeny was largely recapitulated. *P. aloides* was positioned as an external lineage in the V phylogeny, likely reflecting a long-branch effect (**Figure S8**). The concordant species relationships within both U and V clades indicate that the U/V sex chromosomes originated before the diversification of extant Polytrichaceae.

We next estimated SDR age from synonymous divergence (Ks) between U-V gametologs in *P. aloides* (**Figure 2c**). Ks values were very low in the PAR (mean = 6.10-4) but substantially higher in the SDR (mean = 0.22), consistent with strong divergence following recombination suppression. We found no clear evidence for distinct strata of recombination suppression. Similar patterns were observed in *P. commune* (**Figure S10**).

Using the Polytrichaceae crown age distribution (**Figure 2a**, **Figure S2**), the corresponding synonymous divergence distribution (**Figure S3**), and the Ks distribution of SDR gametologs in *P. aloides* (**Figure 2c**; mean = 0.254, SD = 0.088), we estimated the origin of the U/V sex chromosomes at 123.5 Mya (95% interval of the resampled age distribution: 66.1-290.9 Mya). Including Ks values from *P. commune* yielded a similar estimate of 114.7 Mya (95% interval: 66.1–272.3 Mya), despite the use of sexed samples from a more distantly related species. Ks was even higher in *A. angustatum*, for which the U and V chromosomes were assembled separately (mean = 0.53; **Table S8**), suggesting that we may underestimate the age of the SDR.

To test whether the Polytrichaceae U/V chromosomes evolved independently from the sex chromosome system previously characterized in Bryopsida (Carey et al., 2021; Gao et al., 2023), we searched within their SDRs for ancestral Bryopsida U–V gametologs. Of the 16 most divergent U–V gametolog pairs identified in *Ceratodon purpureus*—13 originating at the base of Dicranidae and three predating the split between Bryidae and Dicranidae—only two were sex-linked across all Polytrichaceae species (**Table S9**). These two gametologs showed substantially lower Ks values in *P. aloides* than in Bryopsida (0.01-0.39 *vs* 0.728), consistent with their independent and more recent recruitment into the Polytrichaceae sex chromosomes.

### Polytrichaceae U/V chromosomes show limited gene degeneration

The proportion of genes located in the SDRs of Polytrichaceae sex chromosomes ranged from 0.77% to 1.88% of the total gene content (**Table S4**). The five Polytrichum species had the highest SDR gene content, with an average of 371 genes, compared with 238 genes in *P. aloides* and an average of 204 genes in the other Polytrichaceae species, excluding the polyploid *P. sexangulare*. Consistent with this pattern, reconstruction of SDR gene content along the phylogeny indicated lineage-specific gene gains in Polytrichum (**Figure 2b**). We identified 108 U-V gametolog families inferred to be ancestral to Polytrichaceae, with high Ks values in *P. aloides* consistent with their origin at the base of the U/V chromosomes (**Figure 2b-c**). In addition, many genes were restricted to particular clades or species and showed lower Ks values, indicating more recent recruitment into the SDR. Compared with sex chromosome systems of similar age, Polytrichaceae SDRs retained an unusually high number of genes, consistent with limited gene degeneration (**Figure 3**). Across U/V systems, the proportion of genes located in SDRs was generally below 5% of the total gene content and was particularly low in algae and most liverworts (**Table S4**). The Polytrichaceae SDRs nevertheless contained fewer genes than those of Bryopsida, with *Ceratodon purpureus* being an exception due to its recent autosomal fusions: its sex chromosomes comprise ∼32% of the genome but only ∼11% of its genes (Carey et al., 2021).

**Figure 3.**
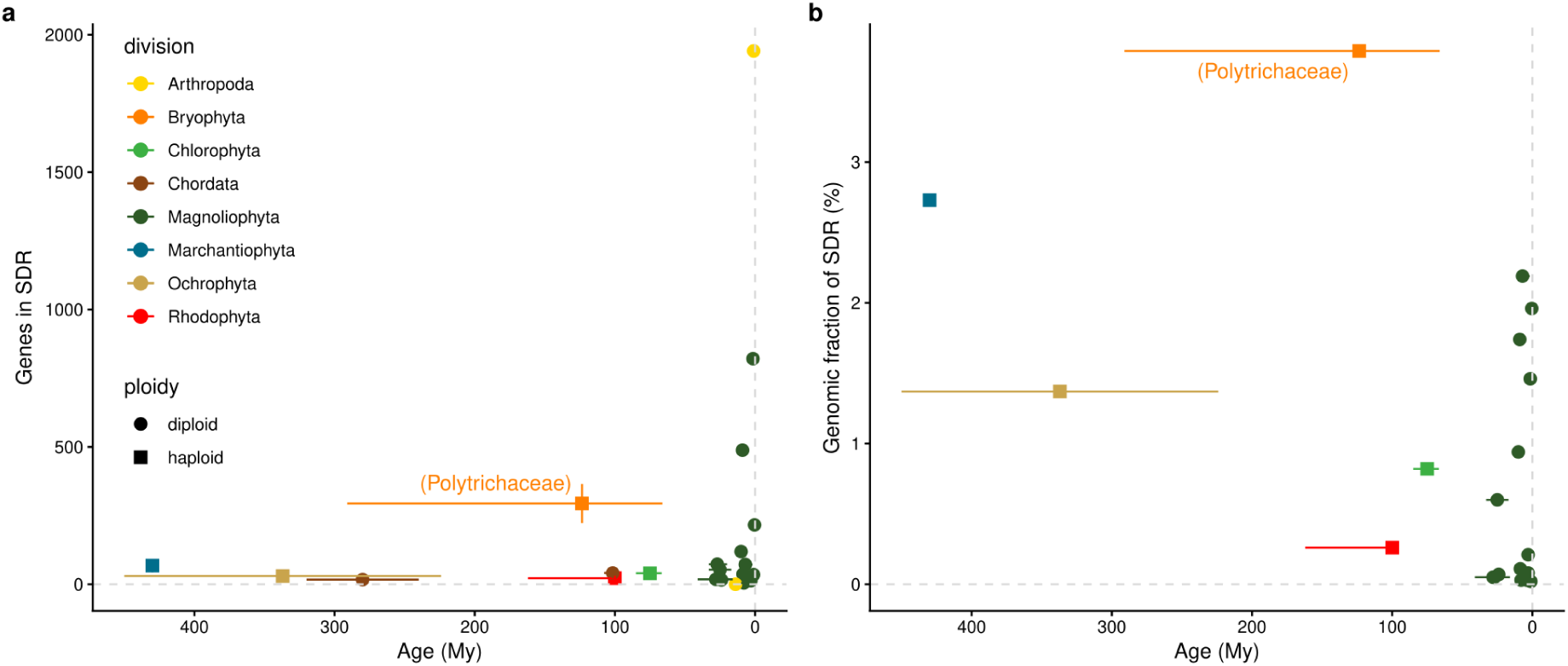
Haploid versus diploid sex chromosome evolution. **(a)** The total number of genes in the SDR and **(b)** the genomic fraction of the SDR are plotted against sex chromosome age for bryophytes (Bryophyta, *n*=1, and Marchantiophyta, *n*=1), algae (Ochrophyta, *n*=1, Rhodophyta, *n*=1, and Chlorophyta, *n*=1), angiosperms (Magnoliophyta, *n*=15), and animals (Arthropoda, *n*=2, and Chordata, *n*=2 for panel (a) only). Angiosperm data were reproduced from Figure 3 of Renner & Müller, 2021, and animal data were compiled from D. Charlesworth, 2021 (see **Table S10** for details). Diploid sex chromosome systems (XY or ZW) are represented by circles and haploid systems (UV) by squares. Horizontal error bars indicate uncertainty in SDR age estimates, whereas vertical bars indicate the standard deviation in SDR gene number when multiple species share the same sex chromosome system. *Ceratodon purpureus* was excluded from the bryophyte dataset because its recent autosomal fusions make it an outlier in this comparison. Colors indicate phyla.

We next assessed the extent of degeneration among genes retained in Polytrichaceae SDRs. Both the fraction of optimal codons (FOP) and GC content at third codon positions (GC3) were lower in SDRs than in PARs and autosomes across Polytrichaceae species (**Figure S11**). The reduction in FOP is consistent with weaker selection on codon usage, whereas the lower GC3 is consistent with the absence of GC-biased gene conversion in the non-recombining SDR. We then used population genomic data from *P. commune* to assess the efficacy of selection on SDR genes (**Figure 4**). Neutral diversity was significantly lower in the SDR (mean πS = 3.10-5) than in the PAR and autosomes (2.10-3), similar to patterns reported in the brown alga Ectocarpus (Avia et al., 2018). This reduced diversity is consistent with strong Hill–Robertson effects associated with suppressed recombination and reduced effective population size. Synonymous divergence measured between *P. commune* and *Polytrichum juniperinum* was also significantly lower in the SDR (mean dS = 0.04) than in the PAR (0.22) and autosomes (0.21). This pattern is consistent with reduced ancestral polymorphism in the SDR, resulting in reduced fixation of neutral mutations along each lineage (B. Charlesworth, 2012). This contrasts with patterns reported in brown algae and *Ceratodon purpureus* (Barrera-Redondo et al., 2025). In Polytrichaceae, both πN/πS and dN/dS were lower in the SDR than in the rest of the genome, although only the difference in dN/dS was significant. This pattern indicates strong selective constraint on genes retained in the SDR. In contrast, the PAR did not differ significantly from autosomes for any of these metrics.

**Figure 4.**
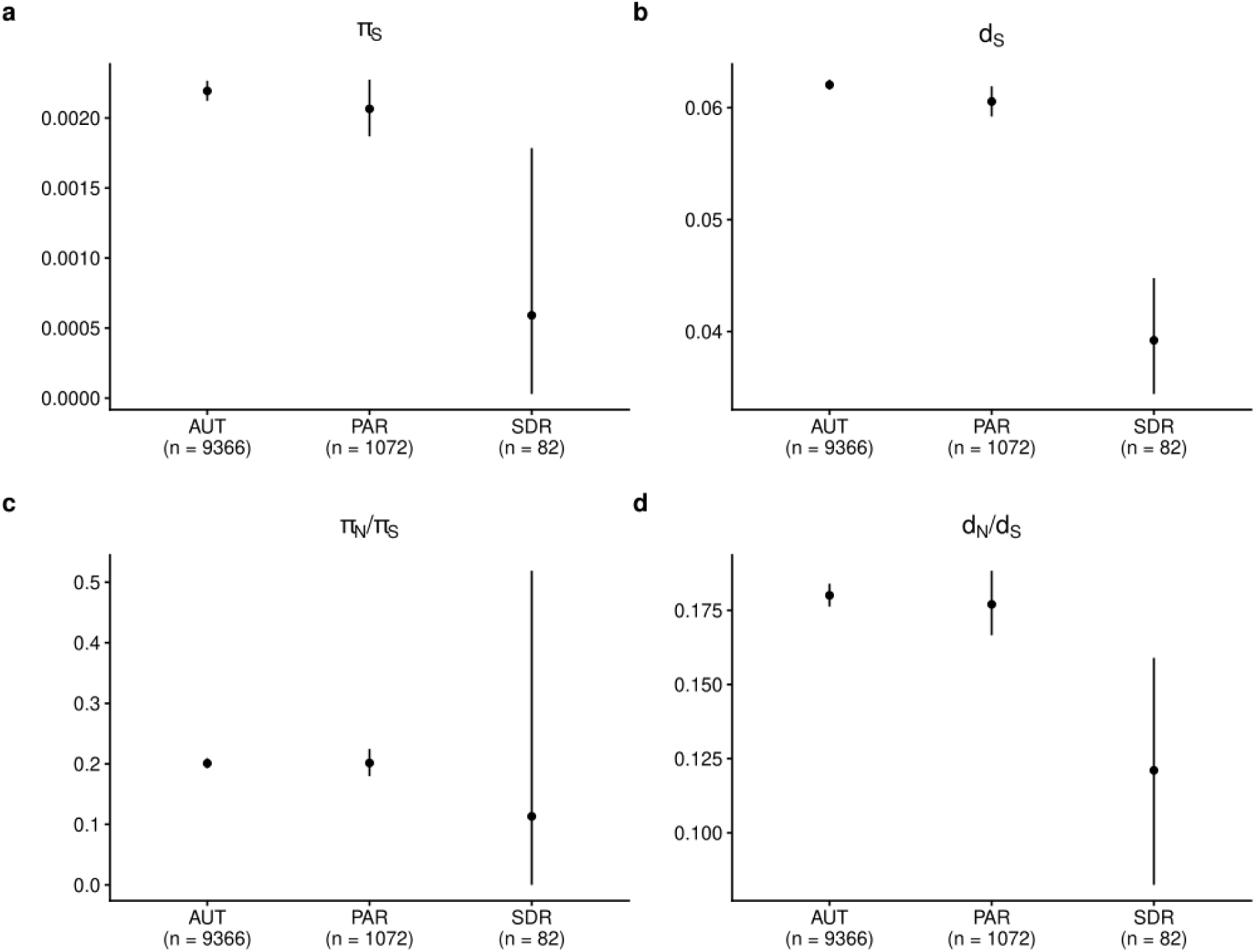
The haploid sex chromosome of *Polytrichum commune* shows signatures consistent with Hill–Robertson effects and limited gene degeneration. **(a)** Synonymous nucleotide diversity (πS). **(b)** Synonymous divergence (dS). **(c)** Non-synonymous to synonymous nucleotide diversity ratio (πN/πS). **(d)** Non-synonymous to synonymous divergence ratio (dN/dS). Dots indicate mean values and vertical bars indicate bootstrap confidence intervals for autosomes (AUT), the pseudo-autosomal region (PAR), and the sex-determining region (SDR). Estimates were calculated from coding sequences of four male *Polytrichum commune var. perigoniale* samples; two male *Polytrichum juniperinum* samples were used as the outgroup for divergence estimates (**Figure S12**). The number of genes included in each estimate is indicated by n.

## Discussion

### Old and independent origins of sex chromosomes in U/V sexual systems

Recent genomic studies have revealed that U/V sex chromosomes can be remarkably ancient in algae and bryophytes, in contrast to the generally younger sex chromosomes of flowering plants (**Table S4**, Renner & Müller (2021)). In bryophytes, ancient U/V systems have been identified in hornworts (Bowman & Levins, 2025; Schafran et al., 2025), liverworts (Iwasaki et al., 2021; Yue et al., 2026) and mosses (Carey et al., 2021; Gao et al., 2023; Healey et al., 2023) with evidence for multiple independent origins among major lineages. Our results identify a third independently evolved U/V system within mosses, in Polytrichaceae, dating to approximately 123 Mya. This age is substantially younger than the U/V chromosomes shared across Bryopsida (>300 Mya), consistent with independent origins rather than inheritance from their common ancestor. In Polytrichaceae, the high Ks divergence between U-and V-linked ancestral gametologs, together with their shared syntenic location across the family, indicates that recombination suppression originated before the radiation of the family. Nevertheless, more recently, lineage-specific gametologs were interspersed with ancestral ones, indicating subsequent gene recruitment within the SDR. Similar mixtures of old and young gametologs have been reported in other bryophyte U/V systems (*Ceratodon purpureus*: Carey, Jenkins, et al., 2021; *Blasia pusilla*: Yue et al., 2026 and *Marchantia polymorpha*: Iwasaki et al., 2021), suggesting that gene recruitment can continue long after the initial establishment of recombination suppression. The repeated emergence and long-term persistence of ancient U/V chromosomes provide a powerful comparative framework for understanding how prolonged recombination suppression shapes sex chromosome evolution in an haploid context.

### Limited SDR expansion

The Polytrichaceae U/V sex chromosomes show a striking SDR–PAR structure that, to our knowledge, has not been described in mosses. Most previously studied bryophyte U/V chromosomes lack a detectable PAR, with the notable exception of the liverwort *Blasia pusilla* and potentially some hornworts (Schafran et al., 2025; Yue et al., 2026). This apparent absence of PARs may partly reflect sampling bias toward highly heteromorphic sex chromosomes that have already lost their PAR. In contrast, the homomorphic U/V chromosomes of Polytrichaceae retain a large PAR spanning most of the sex chromosome, resembling the small SDRs maintained over long evolutionary timescales in several algal U/V systems (**Table S4**). The Polytrichaceae PAR closely resembles autosomes in level of synteny, gene and TE densities, codon usage, nucleotide diversity, and protein evolution, indicating that it has remained largely unaffected by recombination suppression. This raises the possibility that the current PAR originated through a U/V-autosome fusion after an ancestral loss of the original PAR, potentially favored by selection for tighter linkage between sex-linked and sexually antagonistic genes (Otto et al., 2011). However, we found no evidence for such a fusion, which would therefore have to predate the diversification of Polytrichaceae. Alternatively, the ancestral U/V chromosomes may have retained their PAR throughout their evolution. Distinguishing between these scenarios will require broader sampling of closely related Polytrichales and comparative analyses of ancestral chromosome structure.

### A dynamic set of genes

Orthology analysis identified 922 orthogroups containing at least one sex-linked gene across the ten Polytrichaceae species (**Table S12**). Of these, 502 orthogroups (579 genes) were species-specific, indicating extensive gene turnover over evolutionary time. Such turnover may reflect both gene loss from the SDR and recruitment or movement of genes into or out of the non-recombining region. Despite this dynamic gene content, 60 orthogroups comprising 732 genes were retained in the SDRs of all ten species for more than 100 Mya, revealing a conserved ancestral core. Similar turnover has been reported in other ancient U/V systems, including red and brown algae and the liverwort Blasia pusilla (Barrera-Redondo et al., 2025; Lipinska et al., 2025; Yue et al., 2026). Ancestral gene-content reconstruction further revealed substantial lineage-specific gene gains in Polytrichum (**Figure 2b**). Microsynteny analyses indicate that these gains do not systematically originate from the homologous PAR, suggesting that individual gene movement, rather than large-scale SDR expansion, contributes to the dynamic gene content of Polytrichum SDRs (**Figure S13**). The prevalence of autosomal paralogs among U/V-linked genes in *Atrichum angustatum* further supports gene duplication and movement as potential mechanisms underlying SDR gene turnover, as also reported in other U/V systems (Bowman et al., 2017; Lipinska et al., 2025). Together, these results reveal a striking contrast between the long-term conservation of the SDR itself and the rapid turnover of its gene content: a core set of genes has remained sex-linked for more than 100 My, while additional genes have been repeatedly gained and lost across lineages.

### Limited gene degeneration

As in other U/V systems studied to date and consistent with Bull’s prediction (Bull, 1978), neither the U nor V SDR of Polytrichaceae shows extensive gene degeneration, despite long-terminal repeats accumulation, reduced gene density, and increased gene rearrangement relative to autosomes (**Table S4** and **Supp. Results**). The non-recombining SDR nevertheless showed strong signatures of Hill–Robertson effects, including reduced genetic diversity and neutral divergence, as reported in the liverwort Riccia (Fu et al., 2025). Genes in the SDR also showed reduced optimal codon usage, consistent with weaker selection on codon usage under suppressed recombination, as reported in Bryidae and Dicranidae mosses (Carey et al., 2021; Gao et al., 2023) and brown algae (Barrera-Redondo et al., 2025), but contrasting with red algae (Lipinska et al., 2025). Finally, the SDR did not show elevated dN/dS, unlike *Ceratodon purpureus* (Carey et al., 2021), further supporting limited protein-coding degeneration. Thus, despite strong genomic signatures of recombination arrest, Polytrichaceae U/V chromosomes have undergone limited functional degeneration.

### The role of haploid selection

Unlike diploid XY or ZW systems, in which the recombining X or Z provides a direct reference for assessing degeneration of the Y or W, both U and V chromosomes are non-recombining in haploid sexual systems. We therefore used SDR gene content and gametolog retention as proxies for degeneration. Across plants and animals, non-recombining SDR regions of haploid systems tend to retain more genes and account for a larger fraction of the genome than diploid systems of similar age (**Figure 3**). Polytrichaceae was particularly striking, with an unusually large number of SDR genes relative to both chromosome age and genome size. This pattern is consistent with the hypothesis that haploid selection exposes recessive deleterious mutations to selection, limiting their accumulation compared with diploid systems where such mutations can be sheltered in heterozygotes. The proportion of SDR genes retaining a gametolog varies widely among U/V systems (**Table S10**), from ∼27% in *Marchantia polymorpha* and 52% in *Ceratodon purpureus* to ∼53% in *Ectocarpus sp. 7* and ranges from 13% to 55% across the red algae. The 57% observed in *A. angustatum* therefore lies at the upper end of the reported range. In similarly old X/Y or Z/W systems, gametolog retention is below 5% in birds and snakes but reaches 43% in humans (D. Charlesworth, 2021), further suggesting slower degeneration in haploid than diploid sex chromosomes. However, this measure may be inflated by gene gains, such as through autosomal fusions, and is therefore best assessed within clades of similarly aged sex chromosomes and across independent recombination strata, as investigated in Microbotryum mating-type chromosomes (Duhamel et al., 2022).

### SDR expansion: what can haploid systems tell us?

Our results raise a broader question: if haploid selection limits degeneration, what drives SDR expansion in haploid sexual systems? In organisms with an extended haploid phase, sheltering of recessive deleterious mutations can only promote SDR expansion if the relevant genes are predominantly expressed during the diploid phase. In fungi, extended recombination suppression beyond mating-type loci has indeed been reported only in species with predominantly diploid life cycles (Jay et al., 2024). However, long (chromosome-scale) SDRs have evolved repeatedly in bryophytes and green algae despite the expected efficiency of purifying selection during the haploid phase, suggesting that haploid selection alone cannot explain the diversity of U/V chromosome architectures. The mechanisms that suppress recombination may therefore be equally important. In particular, recombination landscapes differ among lineages, and pericentromeric recombination suppression can promote rapid SDR expansion in some systems (Filatov, 2024). By contrast, recombination appears more homogeneous along bryophyte chromosomes (Lang et al., 2018; Szövényi et al., 2021), making this mechanism less obvious in mosses. Sexually antagonistic selection provides another potential explanation. Because U and V chromosomes are permanently confined to females and males, respectively, linkage of sexually antagonistic alleles to the SDR can be strongly favored (Immler & Otto, 2015; Rice, 1987). Consistent with this hypothesis, larger SDRs are associated with greater gamete dimorphism in brown algae (Barrera-Redondo et al., 2025), while the mating-type locus is larger in the oogamous *Volvox* than in the anisogamous *Eudorina* and isogamous *Yamagishiella* (Hamaji et al., 2018). Among the algae studied to date, the absence of a PAR has only been reported in red algae (**Table S4**). Notably, *Gracilaria* spp. are oogamous and their carposporophyte is physically attached to the female gametophyte, potentially resulting in stronger sexual selection, as in bryophytes (Bessho & Sasaki, 2024). Sexual conflict over resource allocation between gametophytes and sporophytes, as well as sexual dimorphism in mosses, provide plausible sources of antagonistic selection that could be tested in this system (see **Supp. Discussion**). More generally, U/V chromosomes offer a powerful comparative framework for disentangling the forces shaping sex chromosome evolution. Their independent origins, permanent haploid exposure, and contrasting rates of SDR expansion and degeneration provide natural experiments for testing how selection, recombination landscapes, and sexual conflict interact to shape non-recombining chromosomal regions.

## Data and code availability statement

Raw sequencing reads have been deposited in the European Nucleotide Archive (ENA) under BioProject accession number PRJEB129078. ENA sample accession numbers are listed in Table S3. All code used to perform the analyses described in this study, together with the corresponding input data and README files, is available in the InDoRES repository at https://doi.org/10.48579/PRO/HITRTQ.

## Authors contributions

Conceptualization: C.F., F.R-G.; Methodology: C.F., F.R-G., X.V.; Investigation: C.F., F.R-G., X.V.; Resources: I.C., A.V., N.C., C.B.; Data Curation: F.R-G. with inputs from M.; Formal analysis: F.R-G. with inputs from M.; Visualization: F.R-G.; Writing: C.F., F.R-G. with inputs from X.V.; Review and editing: all authors; Supervision: C.F., X.V.; Funding acquisition: C.F.

## Acknowledgments

The authors acknowledge the Mésocentre de Calcul Scientifique Intensif de l’Université de Lille for providing access to high-performance computing resources. They also acknowledge the Région Hauts-de-France, the Ministère de l’Enseignement Supérieur et de la Recherche (through the CPER Climibio and CPER Ecrin grants), and the European Regional Development Fund (ERDF) for supporting the molecular biology platform of the Lille group. Finally, we are grateful to the members of the Comité de Suivi Individuel (Nicolas Bierne, Maxime Bonhomme, Nils Cronberg, Juan-Carlos Villarreal, and Pascal Touzet) for their helpful advice.

