## Supplementary text and figures for "Ancient U/V sex chromosomes reveal slow expansion of recombination suppression in Polytrichaceae mosses"

### Supplementary results

#### U and V chromosomes show largely symmetric gene evolution

The presence of both U and V chromosomes in the *Atrichum angustatum* reference genome allowed us to compare their evolutionary trajectories under similar non-recombining conditions and effective size. Of the 449 genes located in the SDR, 193 (43%) were specific to either the U or V chromosome, of which 152 (79%) retained an autosomal copy (**Table S11**). The V chromosome was 21 Mb long and contained 236 genes, whereas the U chromosome was 15 Mb long and contained 213 genes, resulting in similar gene densities. The two chromosomes also had similar numbers of chromosome-specific genes (99 on V and 94 on U), similar transposable element densities and FOP, suggesting broadly symmetric evolution of the male- and female-limited chromosomes.

### Supplementary discussion

#### Sexual antagonism in Polytrichaceae mosses

Although conspicuous sexual antagonism is relatively uncommon in land plants, sexual conflict may arise over resource allocation from the female gametophyte to sporophyte offspring, with maternal and paternal interests favoring different allocation strategies (Haig, 2013; Haig & Wilczek, 2006). The long-lived gametophytes of bryophytes could therefore favor V-linked alleles promoting sporophyte growth and U-linked alleles limiting resource allocation. However, such conflicts are expected to be strongest in outcrossing species that reproduce repeatedly. Polytrichaceae are acrocarpous, with shoots typically producing a single perigonium per year, although the clonal colony is perennial. In males, successive growth points can develop within the perigonium, resulting in a vertical sequence of perigonia that reflects reproductive activity over several consecutive years. Moreover, unlike many other dioicous mosses, Polytrichaceae lack specialized asexual propagules and appear to reproduce asexually infrequently (**Table S3**). However, asexual propagation can also occur through the development of sometimes extensive underground networks of rhizome-like structures. These features may influence the potential for sexual conflict within perennial clonal colonies. Sexual dimorphism could provide another source of sexually antagonistic selection, as several dimorphic traits have been documented in mosses (Carey, Kollar, et al., 2021). However, there is currently no evidence that Polytrichaceae are less sexually dimorphic than other mosses and liverworts. For example, male plants develop a highly specialized ‘splash-cup’ perigonium, with leaves adapted to facilitate the dispersal of male gametes. Spore size has been associated with the degree of sexual dimorphism (Ramsay, 1979), suggesting that sexual differentiation may be reflected in traits expressed as early as the spore stage. This association might reflect reduced nutrient-storage requirements in spores that give rise to dwarf males, which rely on resources provided by the female gametophyte. In line with this, Polytrichaceae, which lack dwarf males, appear to be predominantly isosporous, with no evidence of bimodal spore-size distributions in the species investigated to date (Rocha et al., 2008). However, the absence of such distributions does not preclude sexual dimorphism at later developmental stages. Alternatively, sexual antagonism in Polytrichaceae may be resolved primarily through sex-biased expression of autosomal genes rather than sex chromosomes (McDaniel, 2023).

### Supplementary figures

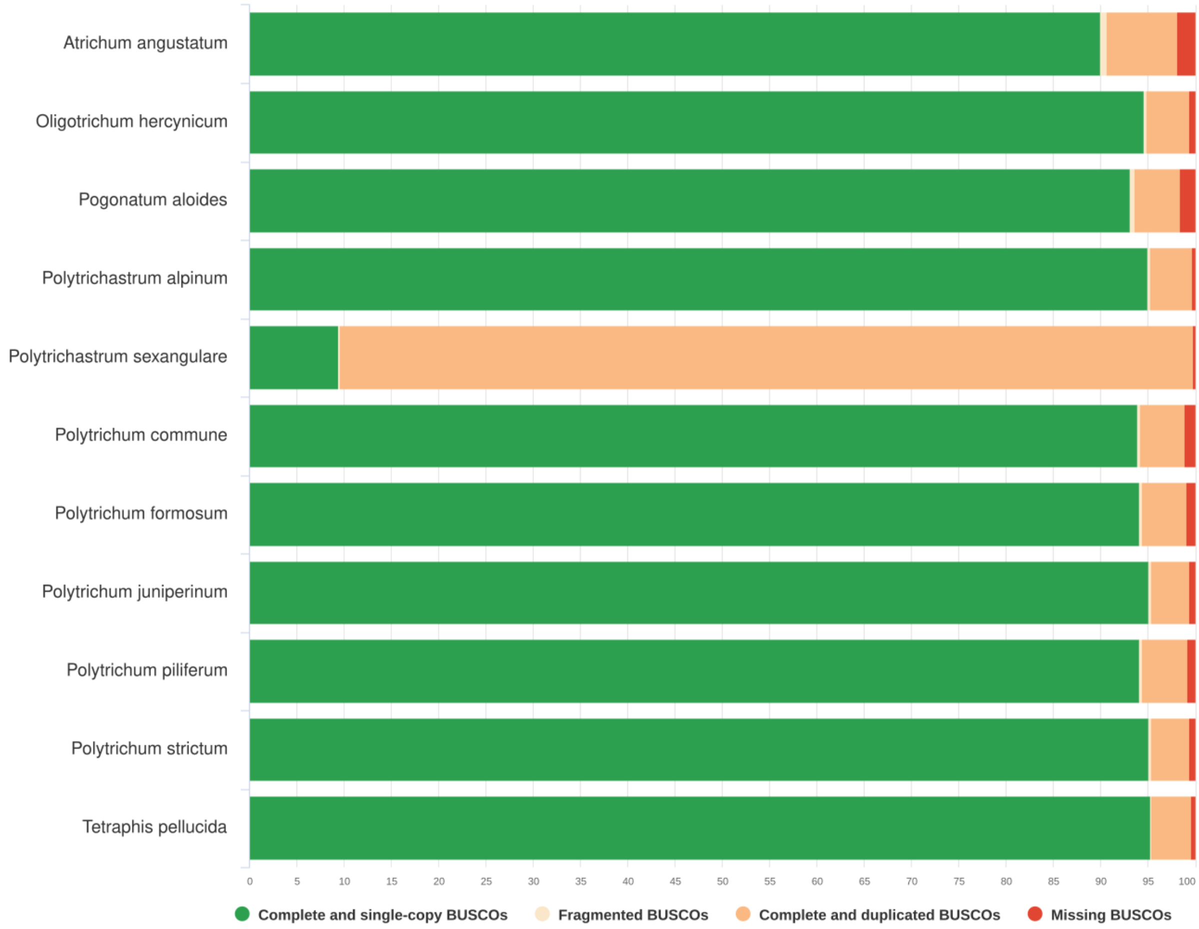


**Figure S1**: BUSCO completeness assessment of genome annotation based on the Galba gene predictions. The bryophyta_odb10 database was used for the assessment. The high proportion of complete and duplicated BUSCOs in *Polytrichastrum sexangulare* is consistent with its recent whole-genome duplication.

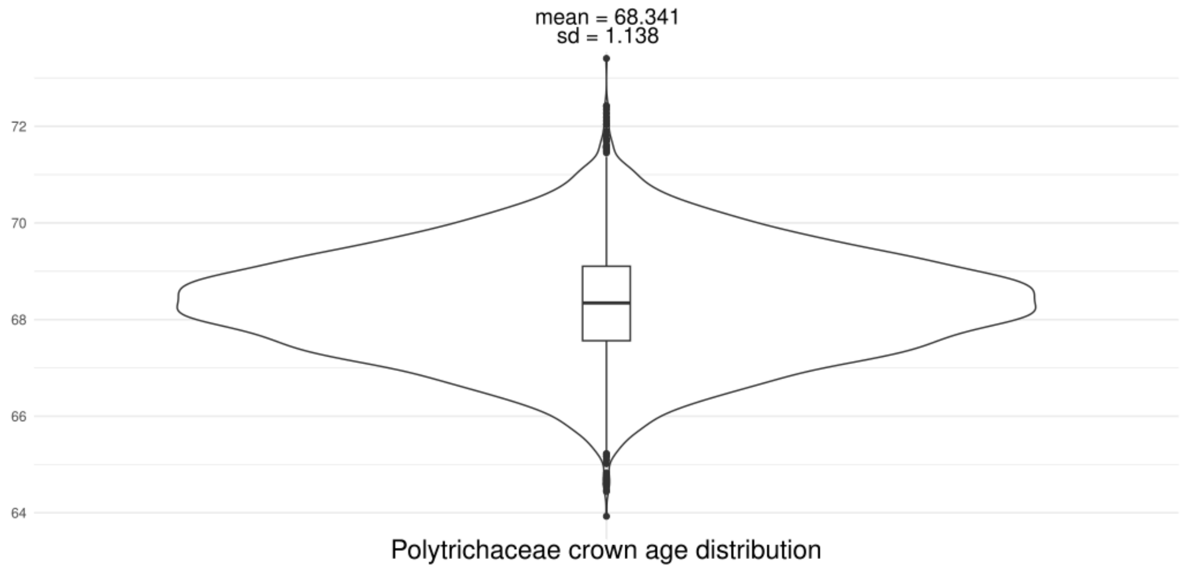


**Figure S2**: Distribution of the Polytrichaceae crown age from 10 million BEAST MCMC iterations, with a 25% burn-in. The mean and standard deviation (sd) are indicated at the top.

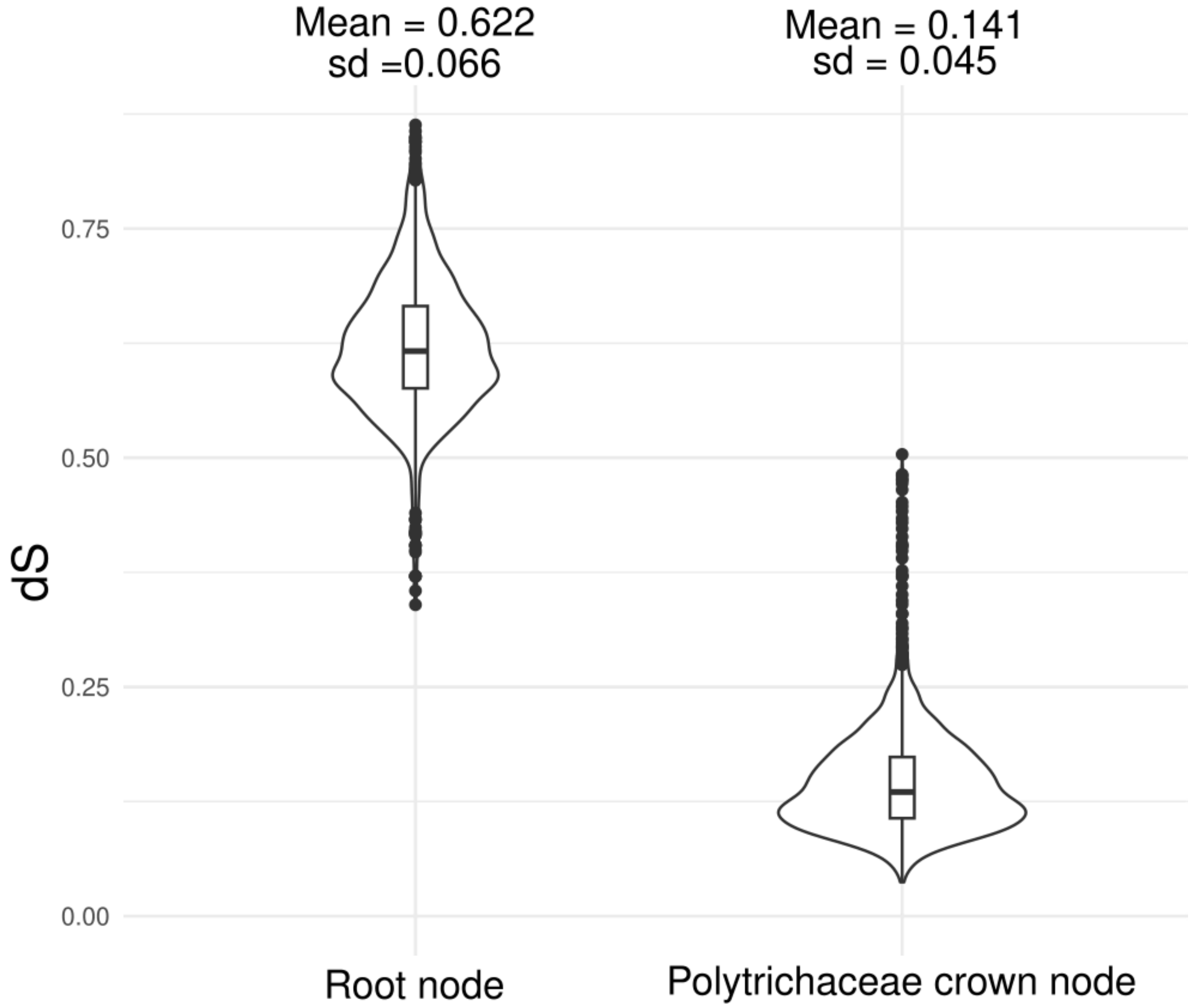


**Figure S3**: Distribution of *dS* values for the 139 BUSCO genes used to reconstruct the species tree. The mean and standard deviation (SD) are indicated at the top. For the root node (left), *dS* was calculated for pairwise comparisons between *Tetraphis pellucida* and each Polytrichaceae species. For the Polytrichaceae crown node (right), *dS* was calculated for pairwise comparisons between species of *Pogonatum* or *Polytrichum* and species of *Atrichum*, *Oligotrichum*, or *Polytrichastrum*.

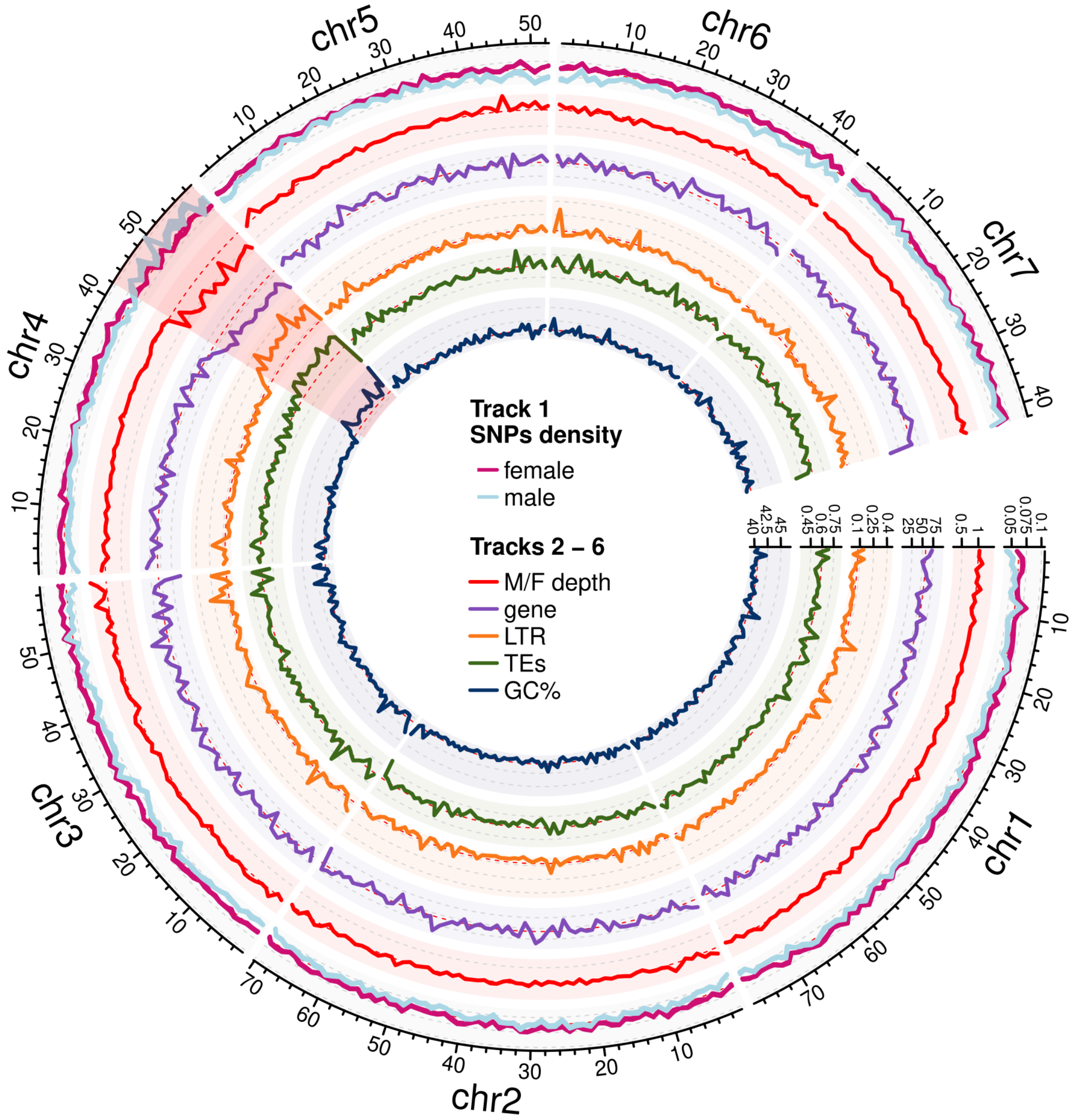


**Figure S4**: Circos plot of genomic features in *Polytrichum commune* shown in 1-Mb sliding windows. From outermost to innermost: (1) SNP density in male (blue) and female (pink) samples; (2) male-to-female sequencing depth ratio; (3) gene density; (4) long-terminal repeat (LTR) density; (5) total transposable element (TE) density; and (6) GC content. The SDR on chromosome 4 is shaded in red. The contrast between males and females is weaker than in *Pogonatum aloides* (Figure 1) because male samples were from the closely related *P. commune var. perigoniale*, whereas female samples were from the more distantly related *Polytrichum formosum*.

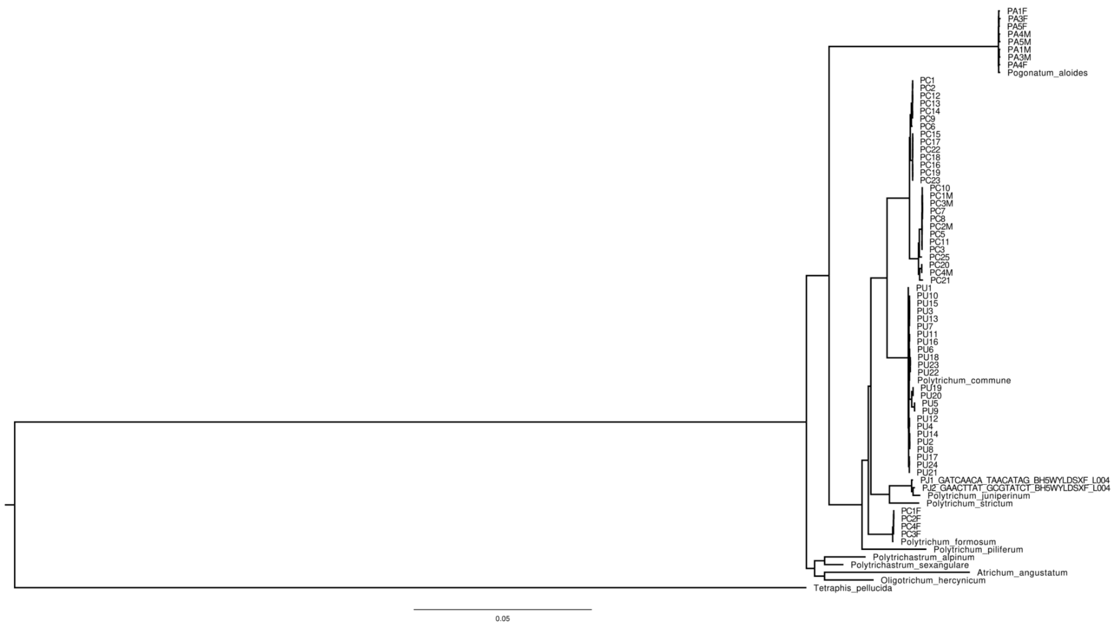


**Figure S5**: Maximum-likelihood phylogenetic tree of the ten Polytrichaceae species studied, inferred with IQ-TREE based on 139 BUSCO genes. Haplotypic sequences from sexed and unsexed samples were also included. Unsexed PC samples (PCx) and male samples (PCxM) were collected from *Polytrichum commune var. perigoniale*. Female samples collected as *P. commune* (PCxF) instead clustered with *Polytrichum formosum*, confirming their misidentification. Unsexed PU samples (PUx) were collected from *Polytrichum commune var. commune*. Male and female samples of *Pogonatum aloides* (PAxM and PAxF) clustered with the reference *P. aloides* sequence. PJ samples, from *Polytrichum juniperinum*, were used as outgroups.

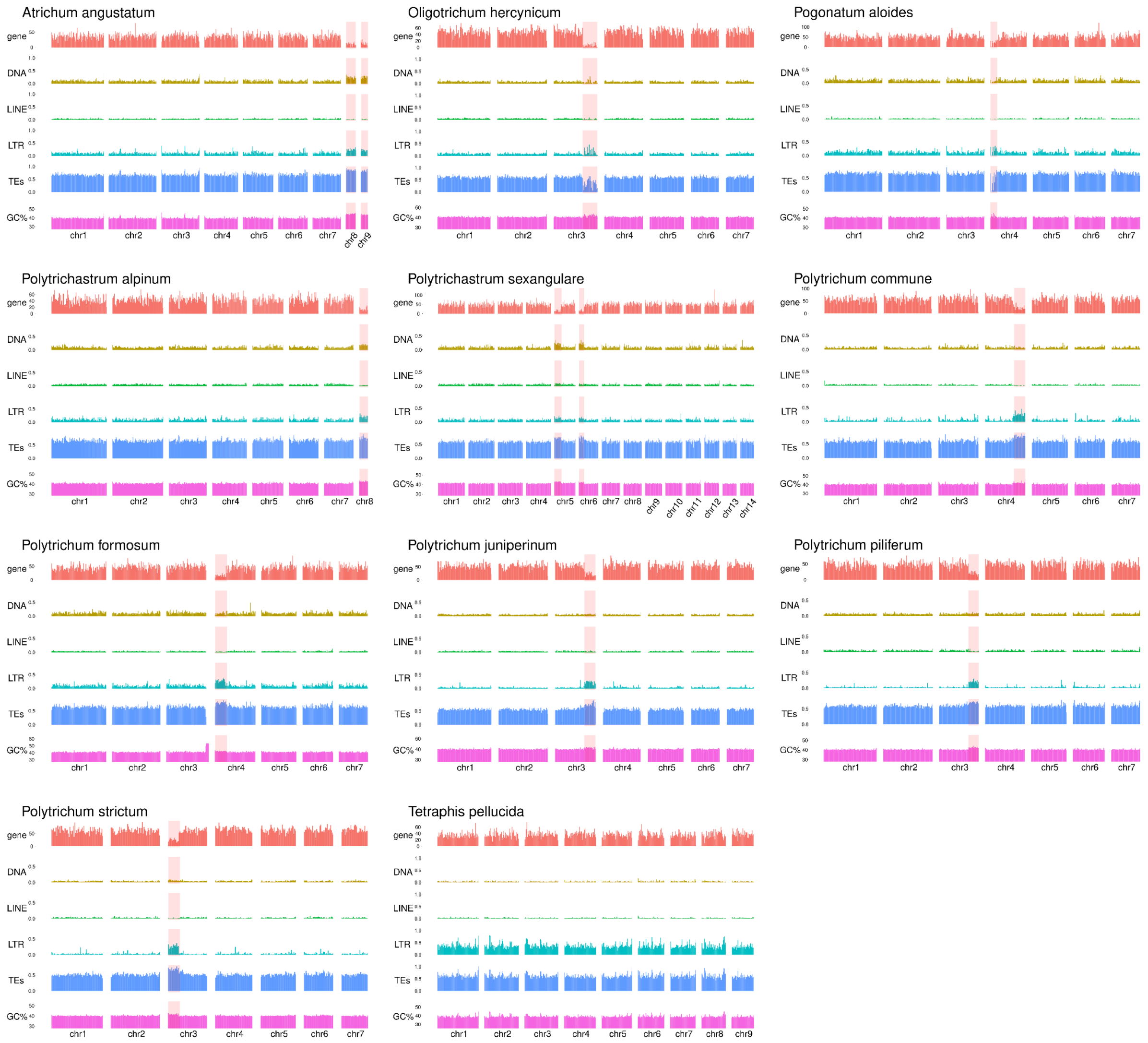


**Figure S6**: Genomic features along the chromosomes of all species studied. From top to bottom: (i) number of genes per Mb; (ii) density of transposable elements (TEs), including DNA transposons, LINEs, LTRs, and total TEs; and (iii) GC content. The SDR is shaded in red.

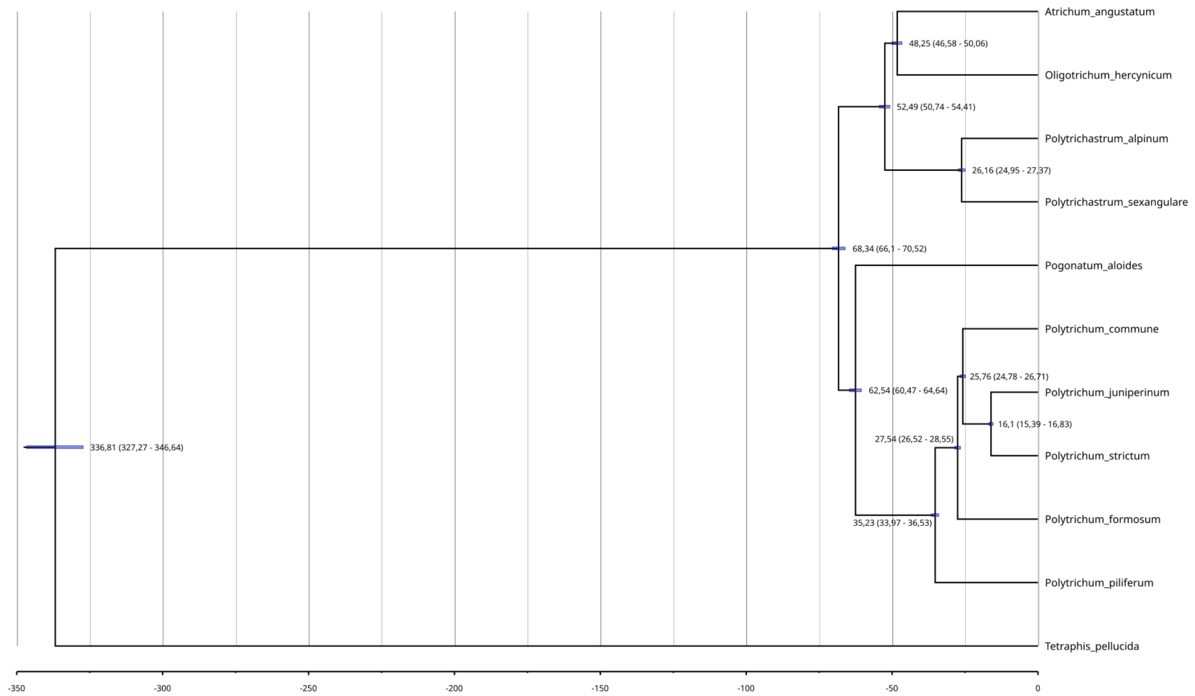


**Figure S7**: BEAST-calibrated phylogeny of the Polytrichaceae species studied, with the mean and 95% highest posterior density (HPD) interval of estimated node ages (Mya). A strict molecular clock was used with a GTR + gamma substitution rate.

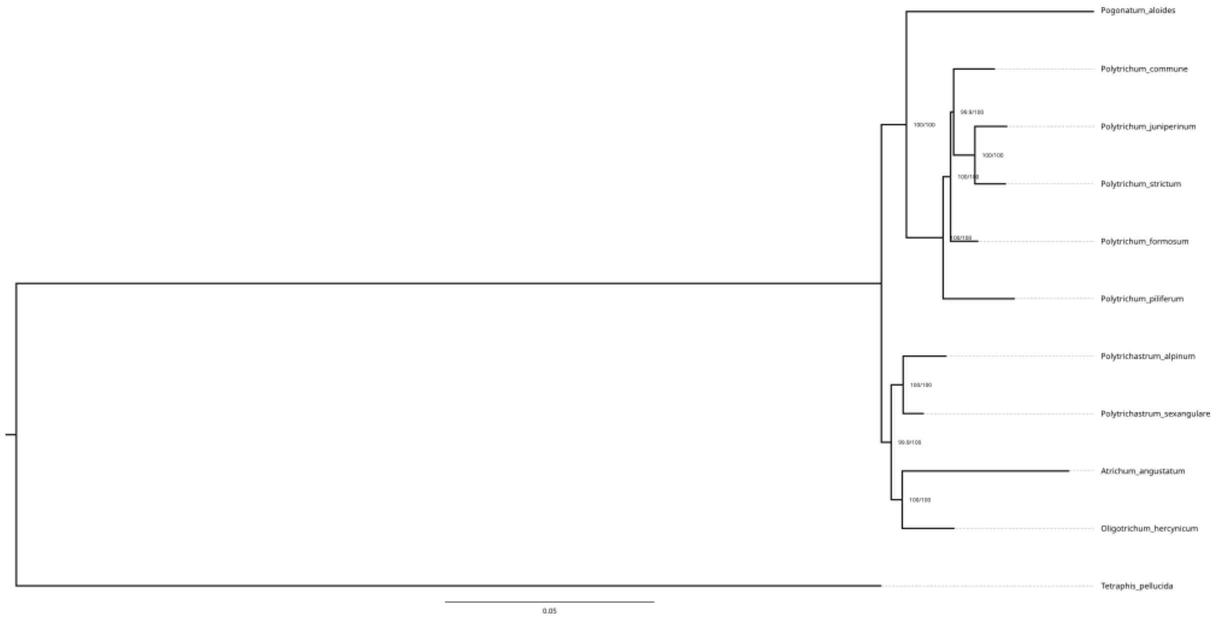


**Figure S8**: Maximum-likelihood phylogenetic tree of the ten Polytrichaceae species studied, inferred with IQ-TREE based on 139 BUSCO genes. Bootstrap support values are indicated at each node.

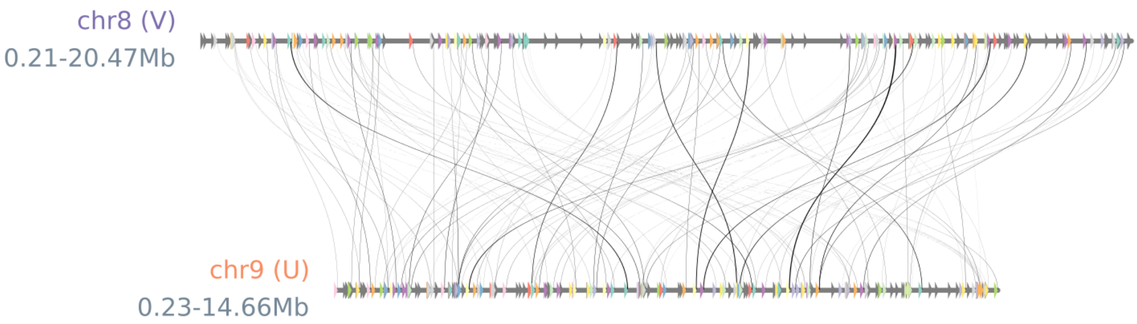


**Figure S9**: Microsynteny relationships between the U (chr9) and V (chr8) sex chromosomes of *Atrichum angustatum*. Each arrow represents a gene, and syntenic genes connected by ribbons are shown in the same color.

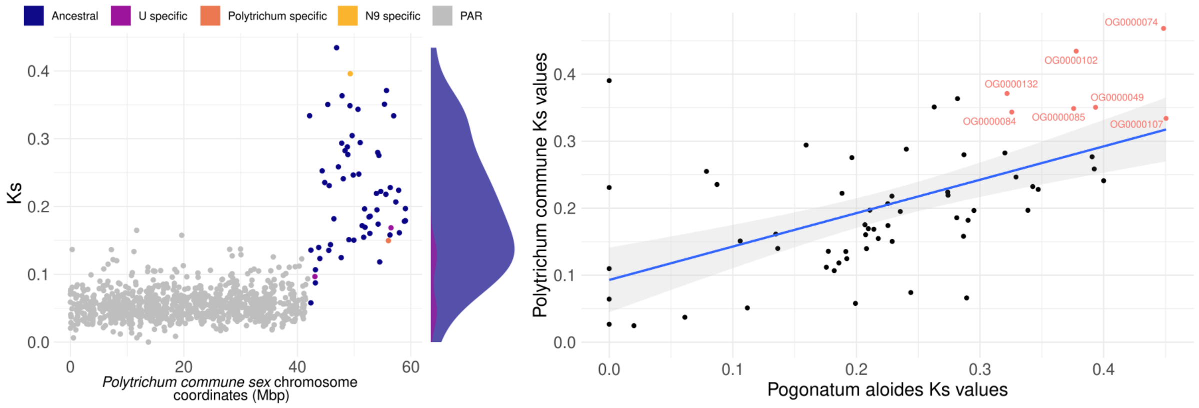


**Figure S10**: Left panel: *Ks* divergence between U- and V-linked gametologs in *Polytrichum commune*, plotted along the sex chromosome. Genes are colored according to their genomic location: PAR (grey), U-specific SDR genes (purple), *Polytrichum*-specific SDR genes (dark orange), *Polytrichum*-specific SDR genes excluding *Polytrichum piliferum* (light orange), and ancestral SDR genes shared across all Polytrichaceae species (blue). Distributions of *Ks* values for ancestral and U-specific gametologs are shown on the right. Right panel: correlation between *Ks* values in *Pogonatum aloides* (**Figure 2c**) and *P. commune* for SDR gametologs, based on orthology inferred with OrthoFinder. The blue line represents the linear regression line, and the shaded area its confidence interval obtained using the *geom_smooth* function in ggplot2 v4.0.3 with method = lm. The seven gametologs with the highest *Ks* values (>0.3) in both species are highlighted in red, with their orthogroup IDs indicated next to each point.

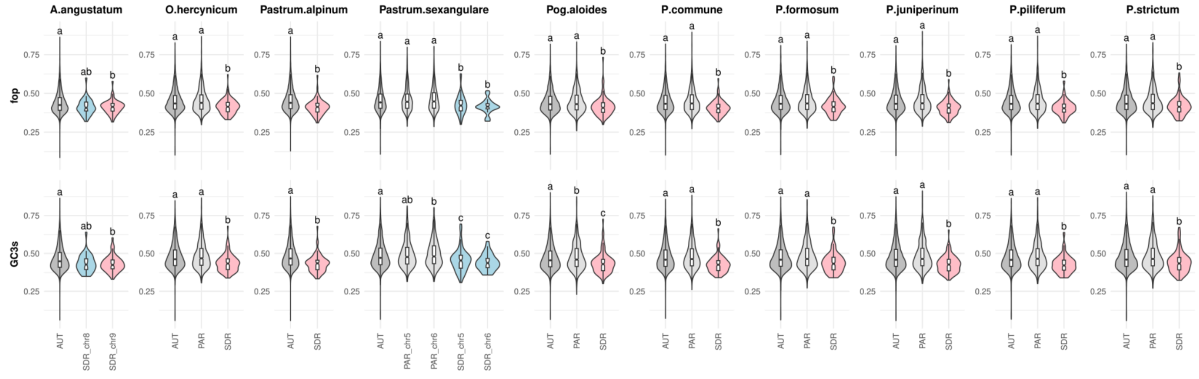


**Figure S11**: Distribution of GC content and codon usage between SDRs (V in lightblue, U in pink), PARs (light grey), and autosomal regions (grey) across the ten Polytrichaceae species studied. fop denotes the fraction of optimal codons. Pairwise Kolmogorov–Smirnov tests were performed. Groups sharing the same letter are not significantly different, whereas different letters indicate significant differences.

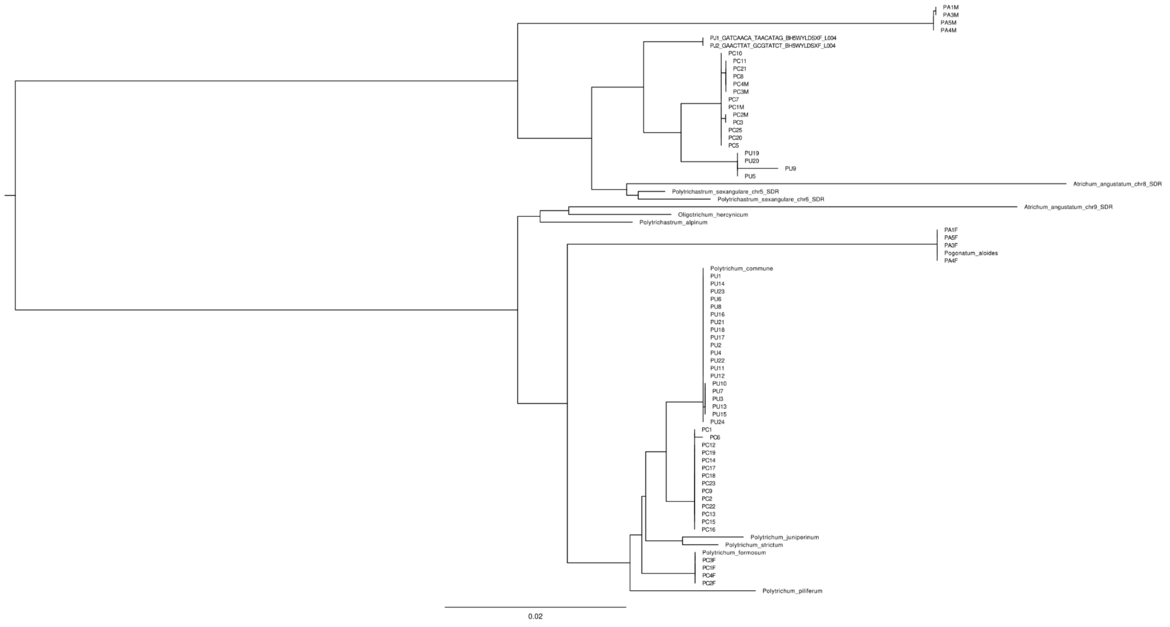


**Figure S12**: Maximum-likelihood phylogenetic tree of the ten Polytrichaceae species studied, inferred with IQ-TREE based on the six orthogroups with the highest *K*s values (>0.3) between gametologs in *Polytrichum commune* and *Pogonatum aloides* (**Table S13**; **Figure S10**). Haplotypic sequences from sexed and unsexed samples were also included, as in **Figure S5**.

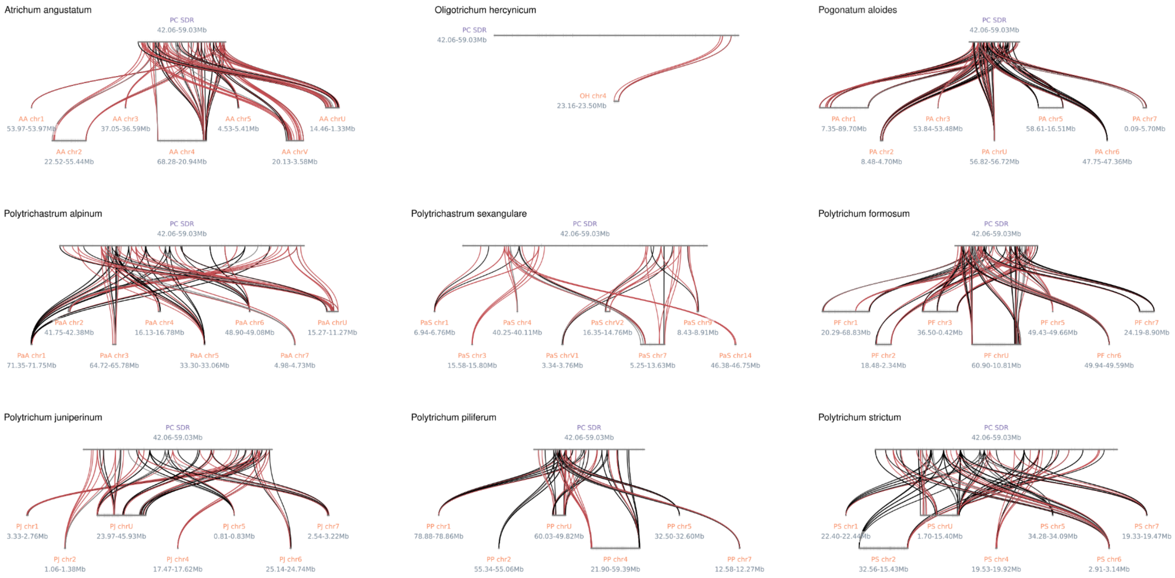


**Figure S13**: Microsynteny relationships between genes acquired in the *Polytrichum commune* U-SDR after *Pogonatum*-*Polytrichum* divergence and genes in each of the other Polytrichaceae species studied. Lines are colored according to the timing of gene gains (see **Figure 2b**). Black lines indicate genes gained in the ancestral lineage of all *Polytrichum* species. Red lines indicate genes gained specifically in *P. commune*. Light grey lines indicate genes gained after the divergence of *Polytrichum piliferum*, whereas grey lines indicate genes gained after the divergence of *Polytrichum formosum*.
